# Serotonergic Axons as Fractional Brownian Motion Paths: Insights into the Self-organization of Regional Densities

**DOI:** 10.1101/2019.12.27.889725

**Authors:** Skirmantas Janušonis, Nils Detering, Ralf Metzler, Thomas Vojta

**Author notes:** **Correspondence:** Dr. Skirmantas Janušonis.

## Abstract

All vertebrate brains contain a dense matrix of thin fibers that release serotonin (5-hydroxytryptamine), a neurotransmitter that modulates a wide range of neural, glial, and vascular processes. Perturbations in the density of this matrix have been associated with a number of mental disorders, including autism and depression, but its self-organization and plasticity remain poorly understood. We introduce a model based on reflected Fractional Brownian Motion (FBM), a rigorously defined stochastic process, and show that it recapitulates some key features of regional serotonergic fiber densities. Specifically, we use supercomputing simulations to model fibers as FBM-paths in two-dimensional brain-like domains and demonstrate that the resultant steady state distributions approximate the fiber distributions in physical brain sections immunostained for the serotonin transporter (a marker for serotonergic axons in the adult brain). We suggest that this framework can support predictive descriptions and manipulations of the serotonergic matrix and that it can be further extended to incorporate the detailed physical properties of the fibers and their environment.

## 1. INTRODUCTION

All cells in vertebrate brains are surrounded by a matrix of highly tortuous fibers that release serotonin (5-hydroxytryptamine, 5-HT), a major neurotransmitter. Presently the self-organization and dynamics of this matrix are not understood beyond neuroanatomical descriptions. Altered densities of serotonergic (serotonin-releasing) fibers have been associated with many mental disorders and conditions, including Autism Spectrum Disorder (Azmitia et al., 2011), Major Depressive Disorder (Numasawa et al., 2017), epilepsy (Maia et al., 2019), and exposure to 3,4-methylenedioxymethamphetamine (MDMA, Ecstasy) (Adori et al., 2011). Predictive computational models can significantly advance this research and support its biomedical applications. Motivated by this potential, we used a stochastic process framework to develop a model of serotonergic fibers and reproduced some key features of their density distribution in the mouse brain.

Serotonergic fibers are axons of neurons whose bodies are located in several brainstem clusters known as the raphe nuclei (Stuesse et al., 1991; Jacobs and Azmitia, 1992; Hornung, 2003). In mammals, these neurons mature early in development. They begin synthesizing serotonin around embryonic day 11-13 in the mouse and rat brains (Lidov and Molliver, 1982; Hendricks et al., 1999; Hawthorne et al., 2010) and around 5 weeks of gestation in the human brain (Sundstrom et al., 1993; Mai and Ashwell, 2004). In the adult mammalian brain, serotonergic axons are unusual in their ability to regenerate, with potential implications for the efforts to restore other axon systems after injury (Hawthorne et al., 2011; Jin et al., 2016; Kajstura et al., 2018). A recent study has shown that they may share this property with other axons in the ascending reticular activating system (Dougherty et al., 2019).

Recent studies have revealed great diversity of serotonergic neurons (Okaty et al., 2019), the functional significance of which is an active area of research (Ren et al., 2018). Paradoxically, transgenic mouse models with no serotonin synthesis in the brain during development have no gross neuroanatomical abnormalities and show only mild behavioral deficits (Mosienko et al., 2015; Pratelli and Pasqualetti, 2019). Serotonergic neurons can release other major neurotransmitters, such as glutamate (Okaty et al., 2019) and perhaps GABA (Stamp and Semba, 1995; Okaty et al., 2019). In the raphe nuclei, serotonergic neurons coexist with many other neurons (Cardozo Pinto et al., 2019; Schneeberger et al., 2019), some of which may participate in stereotyped synaptic arrangements (Soiza-Reilly et al., 2013).

The development of serotonergic fibers is currently conceptualized to proceed in two or three stages: the initial growth in well-defined fiber tracts, followed by extensive arborization and eventual dispersal in “terminal” fields (Lidov and Molliver, 1982; Carrera et al., 2008; Kiyasova and Gaspar, 2011; Jin et al., 2016; Donovan et al., 2019). This orderly sequence is generally consistent with experimental observations at the level of local fiber populations, visualized with tract-tracing techniques or quantified with density measures. It is also theoretically appealing in that it mirrors the development of brain projections that connect two well-defined brain regions (e.g., the lateral geniculate nucleus and the primary visual cortex).

There is little doubt that the initial serotonergic projections form well-defined paths, which we have studied in our own research (Janusonis et al., 2004; Slaten et al., 2010). These paths are well described because they are followed by axon bundles, thus facilitating their visualization in time and space. In contrast, the processes that lead to the formation of regional fiber densities remain poorly understood. Fundamentally, they require a rigorous description of the behavior of single fibers, each one of which has a unique, meandering trajectory. Since these processes are the main focus of the present study, we note several important challenges.

The first detailed morphological description of single serotonergic axons has become available only recently (Gagnon and Parent, 2014). This study has reconstructed a small set of axons originating in the dorsal raphe nucleus and has concluded that they travel through multiple brain regions, rarely branching in some of them and producing profuse arborizations in others. However, true branching points are difficult to distinguish from highly tortuous fiber segments that simply pass each other, especially in bright field microscopy (used in this study). A branching point can be unambiguously demonstrated only by examining an individual fiber at high resolution in all three-dimensions, within the physical section (Pratelli et al., 2017). Even when a confocal system with high-power objectives is used, a branching point can be difficult to distinguish from fibers crossing each other at sub-micrometer distances (Janusonis and Detering, 2019; Janusonis et al., 2019). The extent of local sprouting in regeneration also remains an open problem (Hawthorne et al., 2010; Jin et al., 2016), but current evidence suggests that it may not be significant (Jin et al., 2016). The rapidly developing methods of super-resolution microscopy and tissue expansion are well positioned to provide definitive answers to some of these questions (Janusonis et al., 2019; Wassie et al., 2019).

If serotonergic fiber densities are determined by local arborization, it is unclear how fibers can be restricted to specific “terminal” regions, such as cerebral cortical layers (Linley et al., 2013). Because of their high degree of tortuosity, they are likely to cross over to adjacent areas, suggesting a subtle balance between regional-specific branching and a diffusion-like process. Furthermore, the concept of “terminal region” is ambiguous for the serotonergic axons that typically do not form conventional synapses and can release serotonin at virtually any segment of their trajectory, based on *in vivo* and *in vitro* observations of axon varicosities (Benzekhroufa et al., 2009; Gagnon and Parent, 2014; Quentin et al., 2018). Serotonergic neurons also can release serotonin from the soma, dendrites, and growth cones, effectively making their entire membrane surface active (Ivgy-May et al., 1994; Quentin et al., 2018). It should be noted that serotonergic axons may also form conventional synapses (Papadopoulos et al., 1987), but the extent of this “wiring” transmission (Agnati and Fuxe, 2014) is currently unknown and continues to be debated.

In this study, we model individual serotonergic fibers as paths of a stochastic process that reflects their physical properties and show that regional arborization or other local control is not necessary to arrive at a good approximation of the observed fiber densities. Instead, these densities may strongly depend on the geometry of the brain. Our novel approach may offer insights into the self-organization of serotonergic densities in development and may also explain their stability in adulthood, without assuming the permanence of individual fiber trajectories. Consistent with this hypothesis, a recent study has shown that in the adult mouse brain regenerating serotonergic fibers do not follow the pathways left by degenerated fibers but can still restore the layer-specific densities after cortical injury (Jin et al., 2016).

We focus on Fractional Brownian Motion (FBM), a process first described under this name by Mandelbrot and Van Ness in 1968 (Mandelbrot and Van Ness, 1968). FBM and related stochastic processes have emerged as flexible and theoretically rich models in a variety of physical and biological systems. In particular, they have been used to understand the behavior of fiber-like objects, such as biopolymer chains and chromosomes (Polovnikov et al., 2018; Polovnikov et al., 2019).

FBM extends the normal Brownian motion (BM), which for over a century has served as a standard model to describe simple diffusion and other similar processes (e.g., simple polymer dynamics and stock markets). While BM assumes independence between non-overlapping increments, FBM expands this model by allowing non-zero correlations. The sign and strength of the increment correlation is determined by the Hurst index (*H*), which defines two fundamentally different FBM regimes. If 0 < *H* < ½, two neighboring increments are negatively correlated, which produces highly jittery, “anti-persistent” trajectories (also known as “rough paths”). If ½ < *H* < 1, two neighboring increments are positively correlated, which produces “persistent” trajectories that tend to maintain their current direction. In precise terms, the correlation between two neighboring increments is given by 2^2*H*−1^ − 1 and the mean-square displacement follows the power law ⟨*x*^2^⟩*∼n*^2*H*^, where *n* is the number of performed steps. In this framework, BM becomes a special case of FBM, represented by *H* = ½. According to the scaling of the mean displacement, which can be (in the number or steps) sublinear (for 0 < *H* < ½) or superlinear (for ½ < *H* < 1), FBM can be classified as *subdiffusion* or *superdiffusion*, respectively. Since the trajectories of serotonergic fibers are considerably less jittery than BM and have the tendency to maintain their current direction, one can expect to capture their behavior with superdiffusive FBM. A more careful comparison of FBM sample paths with actual fiber trajectories suggests that the relevant *H* values are likely to lie in a narrower interval, near the upper bound (between 0.7 and 0.9; in contrast, fibers traveling in virtually straight, “ballistic” trajectories can be represented by *H* ≈ 1). In addition, FBM has four convenient properties that make it a natural choice in this context. First, it is a continuous process, which is consistent with the assumption that fibers grow smoothly over time. Second, it has stationary increments, which informally means that its statistical properties do not change as the process evolves (this assumption is reasonable from the biological perspective). Third, it is a self-similar process, which ensures that the estimation of *H* does not depend on the discretization grid of experimental observations. Since time-dependent information is difficult to obtain in growing fibers (e.g., with time-lapse imaging in live animals (Jin et al., 2016)), this property ensures robustness. Fourth, its increments are normally distributed. Assuming that randomness in the fiber trajectory arises from collision-like events in its microenvironment and that each of these events has a small effect on the trajectory, the total effect of these collisions inevitably leads to a normal distribution (by the Central Limit Theorem). Importantly, FBM is the only stochastic process with all of these properties (assuming mean-zero increments).

Although FBM was introduced over a half-century ago, it poses major challenges in theoretical analyses. This is due to the fact that FBM is neither a Markovian process nor a semimartingale (in contrast to BM). A particularly important problem for biological sciences is the behavior of FBM in bounded domains (e.g., in two- or three-dimensional shapes). Reflected BM is well understood (Ito and McKean, 1965), but it is not until very recently that the first description of the properties of reflected FBM (rFBM) have become available, in one-dimensional domains (Wada and Vojta, 2018; Guggenberger et al., 2019; Wada et al., 2019). The present study is the first application of this theoretical framework to serotonergic fiber distributions, on the whole-brain scale. We determine the steady state distributions of superdiffusive rFBM in constrained brain-like domains and show that they approximate neuroanatomical observations.

## 2. MATERIALS AND METHODS

### 2.1. Immunohistochemistry and Imaging

Two adult male mice (C57BL/6J, 8 months of age, The Jackson Laboratory) were deeply anesthetized with a mixture of ketamine (200 mg/kg) and xylazine (20 mg/kg) and perfused transcardially with saline, followed by 4% paraformaldehyde. Their brains were dissected, postfixed in 4% paraformaldehyde overnight at 4°C, cryoprotected in 30% sucrose overnight at 4°C, and sectioned coronally at 40 µm thickness on a freezing microtome. The sections were rinsed in 0.1 M phosphate-buffered saline (PBS, pH 7.2), incubated in 0.3% H_2_O_2_ in PBS for 20 minutes to suppress endogenous peroxidase activity, rinsed in PBS (3 times, 5 minutes each), blocked in 5% normal donkey serum (NDS) and 0.3% Triton X-100 (TX), and incubated in rabbit anti-serotonin transporter IgG (1:5000; ImmunoStar, #24330) with 5% NDS and 0.3% TX in PBS for 2.5 days at 4°C on a shaker. The sections were rinsed in PBS (3 times, 10 minutes each), incubated in biotinylated donkey anti-rabbit IgG (1:1000; Jackson ImmunoResearch, #711-065-152) with 2% NDS and 0.3% TX in PBS, rinsed in PBS (3 times, 10 minutes each), incubated in the avidin-biotin-peroxidase complex (1:100; Vector Laboratories, #PK-6100), rinsed in PBS (3 times, 10 minutes each), developed with 3,3’-diaminobenzidine and H_2_O_2_ using nickel intensification for 5 minutes (Vector Laboratories, #SK-4100), rinsed in PBS (3 times, 5 minutes each), mounted onto gelatin/chromium-subbed slides, allowed to air-dry, and coverslipped with Permount. They were imaged on the Zeiss Axio Imager Z1 system with the following objectives: 1× (NA = 0.025; used for whole sections), 10× (NA = 0.45), 20× (NA = 0.80), and 40× (NA = 1.30, oil). All procedures have been approved by the UCSB Institutional Animal Care and Use Committee.

### 2.2. Preparation of 2D-shapes

The outer border, ventricular spaces, and major white matter tracts of imaged sections were hand-traced in Adobe Illustrator CC by an expert trained in neuroanatomy. To reduce the number of contours, major tracts at the edge of the section were left out of the border contour. The contours were imported into Wolfram Mathematica 12. The outer border contour was split along the median (sagittal) symmetry line. The right side of the contour was smoothed with a moving average and reflected on the left side. The same procedure was used for internal contours symmetric with respect to the median line (e.g., the cerebral aqueduct). Inner contours away from the median line (e.g., the fornix) were smoothed on the right side with a moving average and reflected on the left side. Therefore, the final digitized contours (rational-valued arrays of X- and Y-coordinates) were perfectly bilaterally symmetric, compensating for minor sectioning plane deviations and real (minor) brain asymmetries.

The obtained contours were next reformatted for FBM simulations. They were transformed into *N* × 2 matrices, the rows of which represented consecutive, integer-valued Y-coordinates and the two columns of which represented the leftmost and rightmost X-coordinates of the contour (also integer-valued). To arrive at this format, the original contour coordinates were divided by an integer factor that after rounding produced at least four X-values for each consecutive Y-value, and the minimal and maximal X-values were chosen. Since this procedure effectively reduced the size of the contour, it was enlarged back to its original size by multiplying the integer coordinates by the same factor and filling in the new, empty rows with X-values obtained by linear interpolation between the nearest available X-coordinates. Because this format cannot encode concavities oriented along the Y-axis (e.g., the third ventricle), such concavities were stored as separate inner contours. In the study, all such concavities were centered on the median line, which allowed their easy capture with the maximal X-value left to the line and the minimal X-value right to the line. For the purpose of this study, all inner contours were treated as impenetrable obstacles, irrespective of their physical nature (ventricular spaces, outer border concavities, white matter tracts).

The computer simulations described in detail below were performed on the Frontera supercomputing system (NSF, Texas Advanced Computing Center).

### 2.3. Discrete Reflected FBM

In the simulations, each individual serotonergic fiber was represented as the trajectory of a discrete two-dimensional FBM (Qian, 2003). Consider a random walker starting at position *r*_*0*_ and moving according to the recursion relation *r*_*n*+1_ = *r*_*n*_ + *ξ*_*n*_, where *ξ*_*n*_ is a two-component fractional Gaussian noise. This means that the *x* and *y* components of *ξ*_*n*_ are Gaussian random numbers with zero average and variance *σ*^2^, and that each component features long-range correlations between the steps (but the *x* and *y* motion are independent of each other). The corresponding covariance function is given by 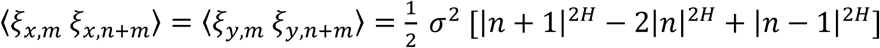, where *H* is the Hurst index. The Fourier-filtering method (Makse et al., 1996) was employed to generate these long-range correlated random numbers on the computer.

If the random walker encounters a boundary (i.e., an outer or inner contour), it is reflected. This reflection can be implemented in several different ways, including a repulsive potential force or a simple reflection condition that either simply prevents the trajectory from entering the forbidden region or mirrors it at the reflecting wall. Extensive test calculations (Vojta et al., unpublished) have demonstrated that the choice of reflection condition does not affect the resulting probability density for rFBM outside a narrow region (just a few steps wide) at the boundary. The following simulations thus employed the condition that a step that would lead the random walker into the forbidden region was simply not carried out.

### 2.4. Reflected FBM-paths in Brain-like Shapes

The goal of the FBM simulations in brain-like shapes was to model the two-dimensional distributions of serotonergic fiber densities at different rostro-caudal levels. Most simulations were performed with *H* = 0.8, but values between *H* = 0.3 and 0.9 were also studied for comparison. Each dataset consisted of 960 individual fibers (FBM trajectories) starting from random positions inside the shape. Each trajectory had 2^22^ ≈ 4 million steps of size (standard deviation) σ = 1.29 μm. The local density (*d*_*s*_) was determined by counting the total number of random walk segments inside each cell (of linear size 12.9 μm) of a square grid covering the shape. The trajectories were sufficiently long for the relative densities to reach a steady state (i.e., they did not change if the trajectory lengths were increased further).

For comparison between the simulated fiber densities and the densities observed in the actual (immunostained) sections, the simulated densities were scaled to “optical densities” by the (Beer-Lambert law-like) transformation *d*_*o*_ = 1 − *exp(*−*kd*_*s*_), where the attenuation parameter *k* was chosen such that the mean pixel value in the simulated section matched the mean pixel value in the image of the immunostained section. This transformation removed the dependence on the arbitrarily chosen number and length of the FBM trajectories, and it also realistically constrained density values to a finite interval.

### 2.5. Reflected FMB-paths in a Ring with Varying Curvature

In order to gain general insights into how the density of FBM-trajectories varies as a function of *H* and the contour curvature, we investigated a ring-like shape bounded by the contours *R*_*outer*_ = 100(1 + 0.5*cos*^2^(4*φ*)) and *R*_*inner*_ = 50 (defined in polar coordinates, where *φ* ∈ [0,2*π*)). This shape was intended to represent the cross section of a highly abstracted vertebrate brain. Vertebrate brains develop from the neural tube and remain topologically tube-like in adulthood, where the “hollow” inside of the tube is the ventricular space filled with the cerebrospinal fluid (CSF).

These simulations were performed with *H* = 0.3, 0.5, and 0.8. Each data set consisted of 100 individual fibers (FBM trajectories), starting from random positions inside the shape. Each trajectory had 2^20^ ≈ 1 million steps of size (standard deviation) σ = 0.4. In the subdiffusive case (*H* = 0.3), it was necessary to increase the step size to 1 to ensure that the simulations reached the stationary regime. As before, the local fiber density (*d*_*s*_) was determined by counting the total number of random walk segments inside each cell (of linear size 1) of a square grid covering the shape.

### 2.6. Simulation of Reflected FBM-paths in a 2D-disk with Crowding

The brain tissue is a highly crowded space, but little quantitative information is available about the extent and geometry of the extracellular space in different brain regions (Hrabetova et al., 2018). In order to assess the sensitivity of our results to crowding effects, we performed FBM simulations in a large disk of radius *R* = 100, filled with 1013 small disk-shaped obstacles of radius *R*_*obs*_ = 2. The obstacles were located within a circle of radius 90, leaving an outer ring of width 10 unoccupied. This shape was intended to represent a highly abstracted neocortical region, with the empty outer rim representing cortical layer I. In adulthood, this layer is nearly devoid of neuron somata (but contains many dendrites and axons that for simplicity were ignored in the simulation).

The simulations were performed with *H* = 0.8. Each dataset consisted of 192 individual fibers (FBM trajectories) starting from random positions inside the shape. Each trajectory had 2^25^ ≈ 34 million steps of size (standard deviation) σ = 0.4. As before, the local fiber density (*d*_*s*_) was determined by counting the total number of random walk segments inside each cell (of linear size 0.5) of a square grid covering the shape.

## 3. RESULTS

Serotonergic fiber densities are typically described with regard to specific neuroanatomical brain regions, with the assumption that they reflect the region’s functional demands and are supported by local biological factors. Our immunolabeling results (Fig. 1) are consistent with the previously reported density maps in the rat and hamster brains (Steinbusch, 1981; Morin and Meyer-Bernstein, 1999). In particular, they show high fiber densities in the superficial layers of the cerebral cortex, consistent with observations in the rat and ferret (Voigt and de Lima, 1991a; Linley et al., 2013).

**Figure 1.**
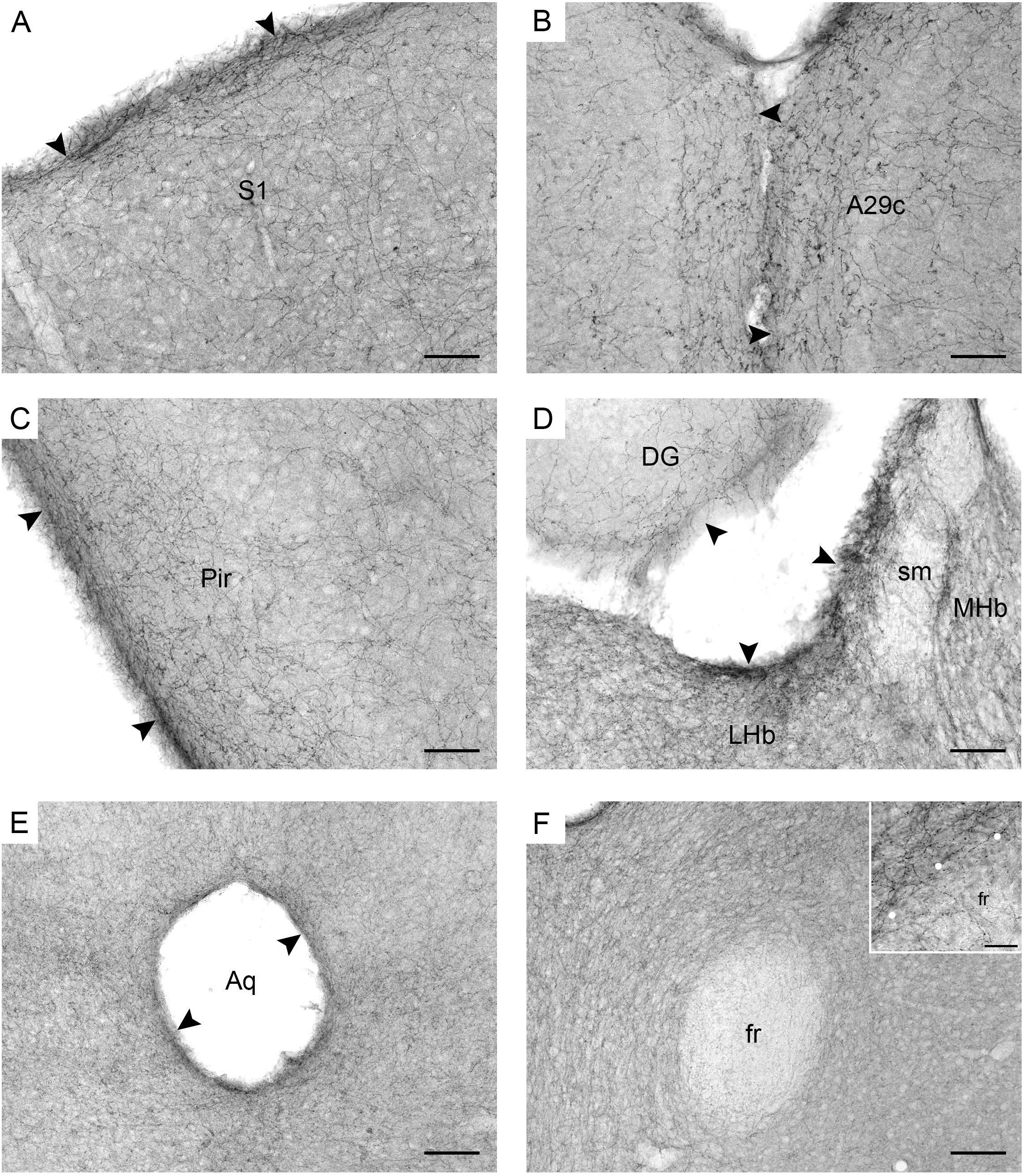
Serotonergic fibers visualized with SERT-immunohistochemistry in coronal sections of the mouse brain. (A) The primary somatosensory cortex (S1). (B) The cingulate cortex, area 29c (A29c). (C) The piriform cortex (Pir). (D) The habenula (MHb, medial habenula; LHb, lateral labenula; sm, stria medullaris) and the dentate gyrus (DG) of the hippocampus. (E) The periaqueductal region in the mesencephalon (Aq, cerebral aqueduct). (F) The region around the fasciculus retroflexus (fr) in the diencephalon. The inset is a high-magnification image of the border (marked with circles) between the fasciculus retroflexus and the surrounding brain tissue. The arrowheads indicate increased fiber densities. Scale bars = 50 µm in A-D, 100 µm in E-F, and 20 µm in the inset of F.

In contrast to previous interpretations, we suggest that these data indicate a *general* tendency of serotonergic fibers to accumulate near the borders of neural tissue (at the pial or ependymal surfaces) (Fig. 1A-E). This observation has been previously made in some brain regions (e.g., in the hamster thalamus (Morin and Meyer-Bernstein, 1999)) but to our knowledge has never been extended to the entire brain. We next show that this tendency is consistent with the behavior of rFBM in the superdiffusion regime.

Supercomputing FBM simulations (with *H* = 0.8) were performed in four two-dimensional shapes that closely approximated the shape of actual coronal sections (Figs. 2-5). Since major white matter tracts may act as forbidden regions (Fig. 1F), we included them as “obstacles” in the simulations. The simulated and actual fiber densities showed similar increases at the outer brain border, irrespective of the rostro-caudal level. A similar trend was observed around the ventricles. Interestingly, high simulated fiber densities were obtained in some neuroanatomically-defined brain regions, such as the lateral geniculate nucleus and the hypothalamus (Fig. 3), even though these regions were not specifically modeled and the increase was induced purely by the contour geometry (Figs. 3-4). It should be noted that both of these regions have a convex border with a relatively high curvature. We investigated this potential association more systematically (Fig. 6), which further supported this conjecture. It leads to verifiable predictions in comparative neuroanatomy, where differences in the serotonergic fiber densities across mammalian species may be caused, at least in part, by differences in the brain shapes (despite the highly similar neuroanatomical plans).

**Figure 2.**
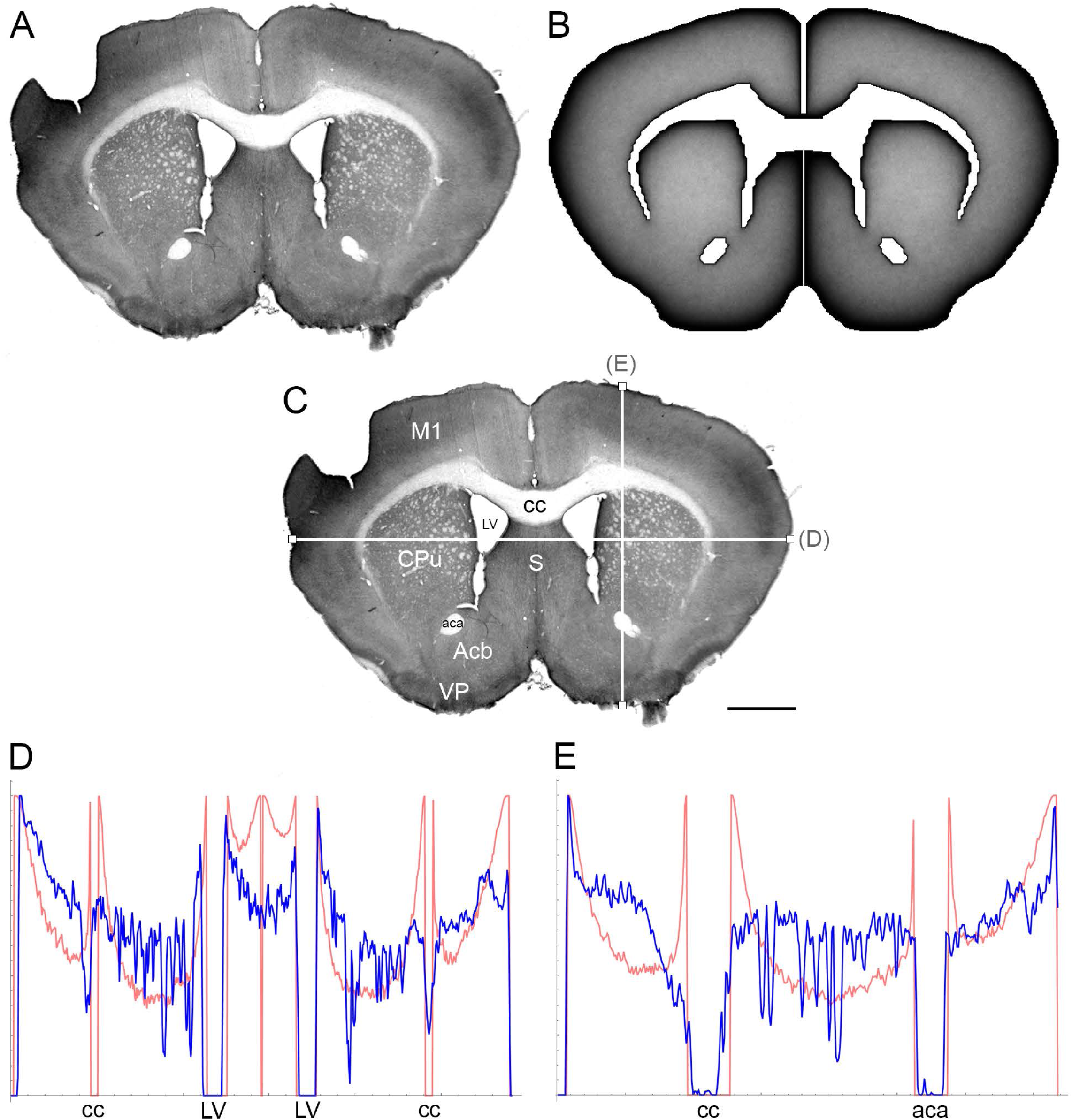
(A) The density of serotonergic fibers (immunostained for SERT) in a coronal section of the mouse telencephalon. (B) The equilibrium density of simulated fibers performing FBM-walks in the same 2D-shape (*H* = 0.8; 960 fibers). In both (A) and (B), darker regions represent higher densities. (C) The main neuroanatomical structures and two density cuts, plotted in (D) and (E). The end points of the plotted segments are marked with small squares. Acb, nucleus accumbens; aca, anterior commissure; cc, corpus callosum; CPu, caudate/putamen; LV, lateral ventricle; M1, primary motor cortex; S, septum; VP, ventral pallidum. Scale bar = 1 mm. In (D) and (E), the experimental and simulated densities are shown in blue and red, respectively. The simulated densities have been transformed to “optical densities,” as described in Materials and Methods (the Y-axis ranges from 0 to 1).

**Figure 3.**
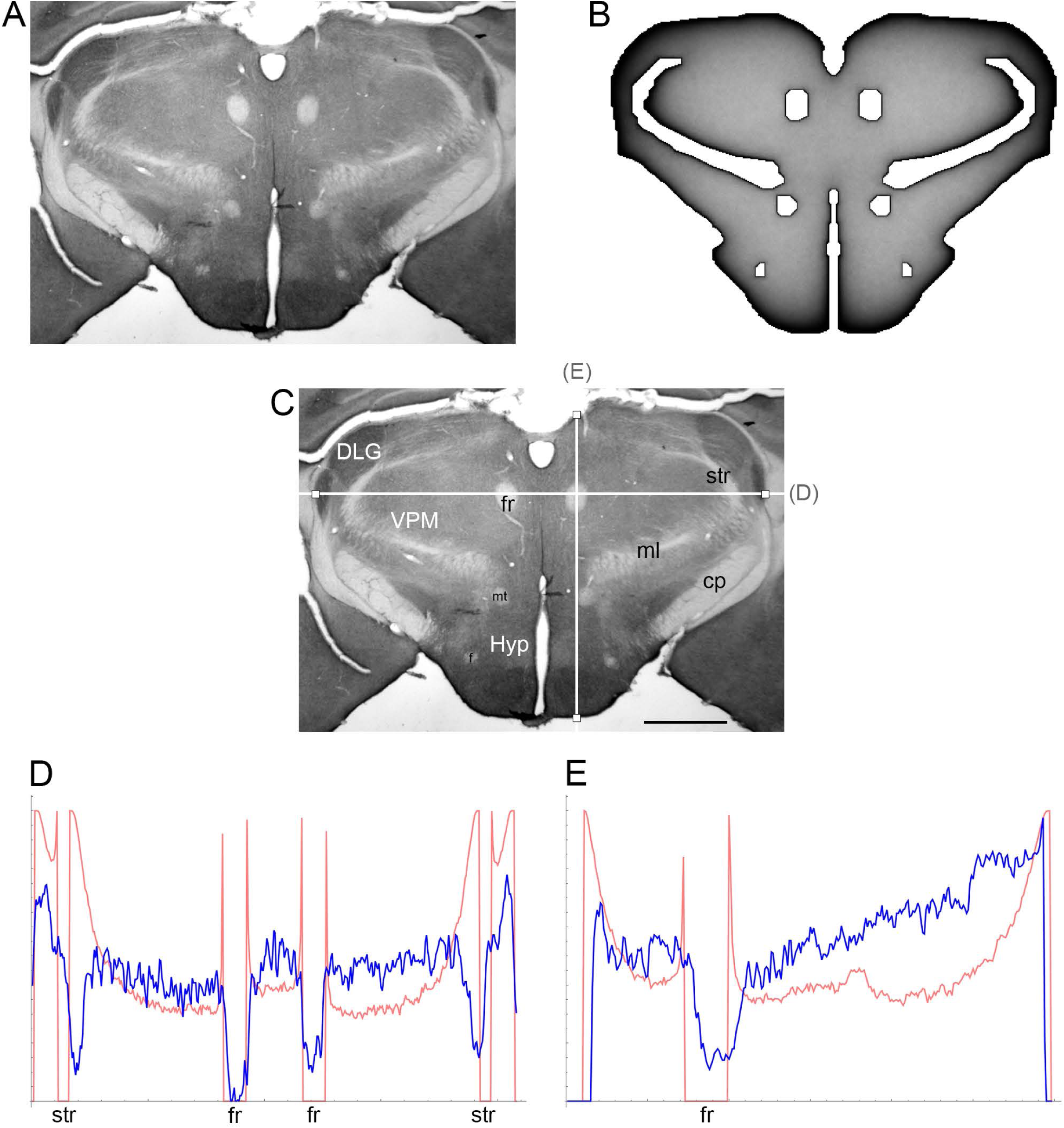
(A) The density of serotonergic fibers (immunostained for SERT) in a coronal section of the mouse rostral diencephalon. (B) The equilibrium density of simulated fibers performing FBM-walks in the same 2D-shape (*H* = 0.8; 960 fibers). In both (A) and (B), darker regions represent higher densities. (C) The main neuroanatomical structures and two density cuts, plotted in (D) and (E). The end points of the plotted segments are marked with small squares. DLG, dorsal lateral geniculate nucleus; cp, cerebral peduncle; f, fornix; fr, fasciculus retroflexus; Hyp, hypothalamus; ml, medial lemniscus; mt, mammillothalamic tract; str, superior thalamic radiation; VPM, ventral posteromedial nucleus. Scale bar = 1 mm. In (D) and (E), the experimental and simulated densities are shown in blue and red, respectively. The simulated densities have been transformed to “optical densities,” as described in Materials and Methods (the Y-axis ranges from 0 to 1).

**Figure 4.**
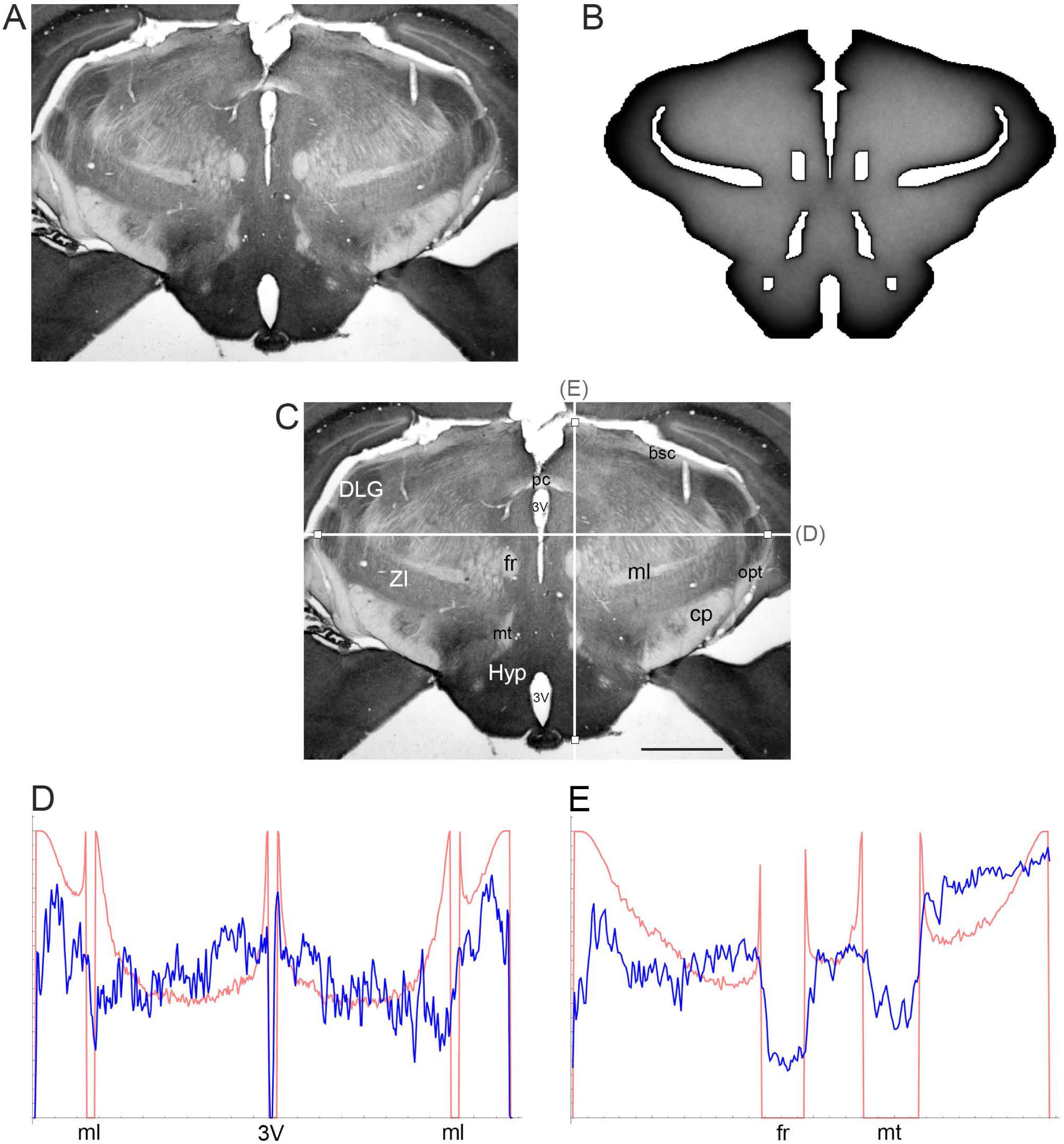
(A) The density of serotonergic fibers (immunostained for SERT) in a coronal section of the mouse caudal diencephalon. (B) The equilibrium density of simulated fibers performing FBM-walks in the same 2D-shape (*H* = 0.8; 960 fibers). In both (A) and (B), darker regions represent higher densities. (C) The main neuroanatomical structures and two density cuts, plotted in (D) and (E). The end points of the plotted segments are marked with small squares. 3V, third ventricle; bsc, brachium of the superior colliculus; cp, cerebral peduncle; DLG, dorsal lateral geniculate nucleus; fr, fasciculus retroflexus; Hyp, hypothalamus; ml, medial lemniscus; mt, mammillothalamic tract; opt, optic tract; pc, posterior commissure; ZI, zona incerta. Scale bar = 1 mm. In (D) and (E), the experimental and simulated densities are shown in blue and red, respectively. The simulated densities have been transformed to “optical densities,” as described in Materials and Methods (the Y-axis ranges from 0 to 1).

**Figure 5.**
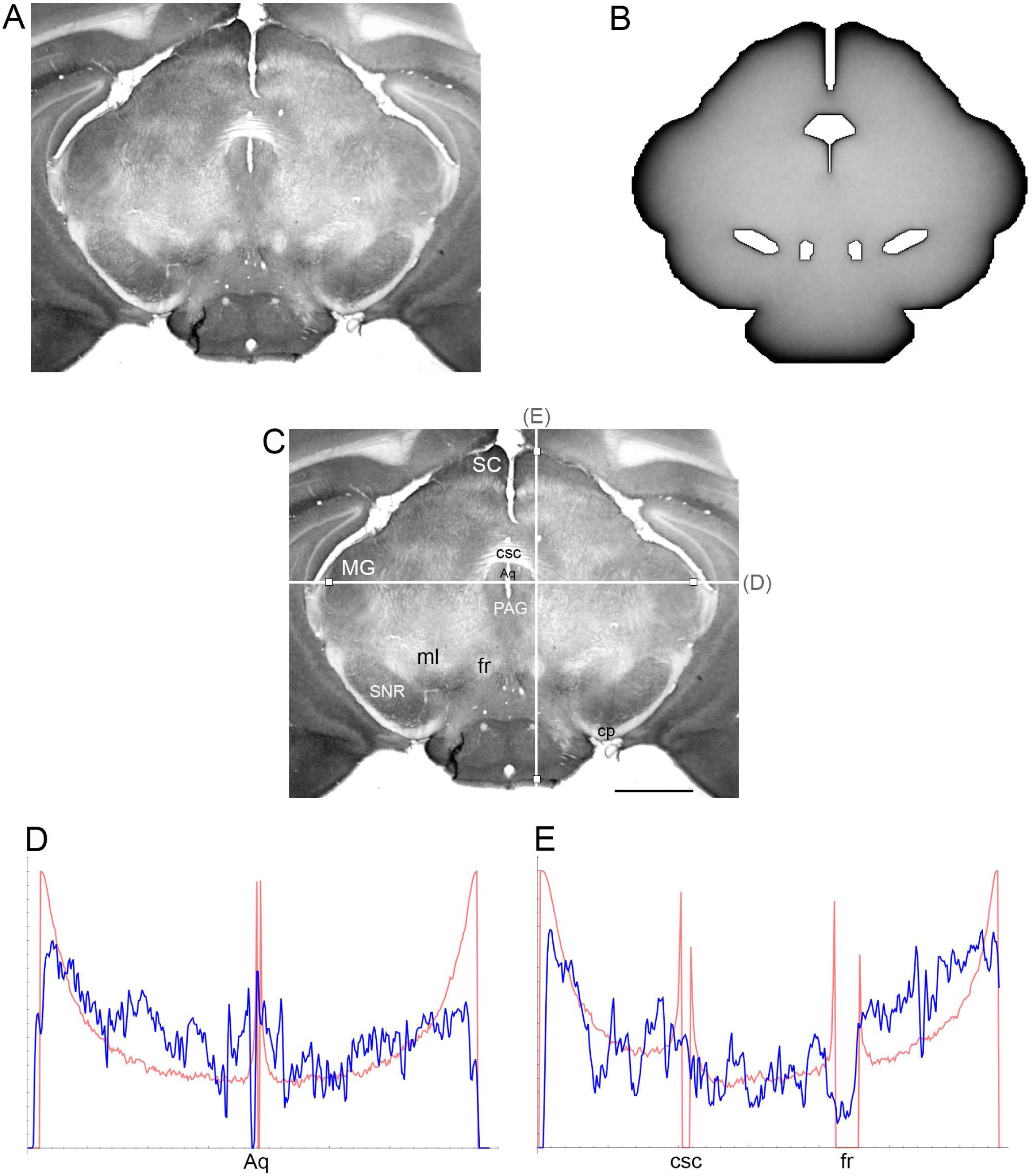
(A) The density of serotonergic fibers (immunostained for SERT) in a coronal section of the mouse mesencephalon. (B) The equilibrium density of simulated fibers performing FBM-walks in the same 2D-shape (*H* = 0.8; 960 fibers). In both (A) and (B), darker regions represent higher densities. (C) The main neuroanatomical structures and two density cuts, plotted in (D) and (E). The end points of the plotted segments are marked with small squares. Aq, aqueduct; cp, cerebral peduncle; csc, commissure of the superior colliculus; fr, fasciculus retroflexus; MG, medial geniculate nucleus; ml, medial lemniscus; PAG, periaqueductal gray; SC, superior colliculus; SNR, substantia nigra pars reticulata. Scale bar = 1 mm. In (D) and (E), the experimental and simulated densities are shown in blue and red, respectively. The simulated densities have been transformed to “optical” densities, as described in Materials and Methods (the Y-axis ranges from 0 to 1).

**Figure 6.**
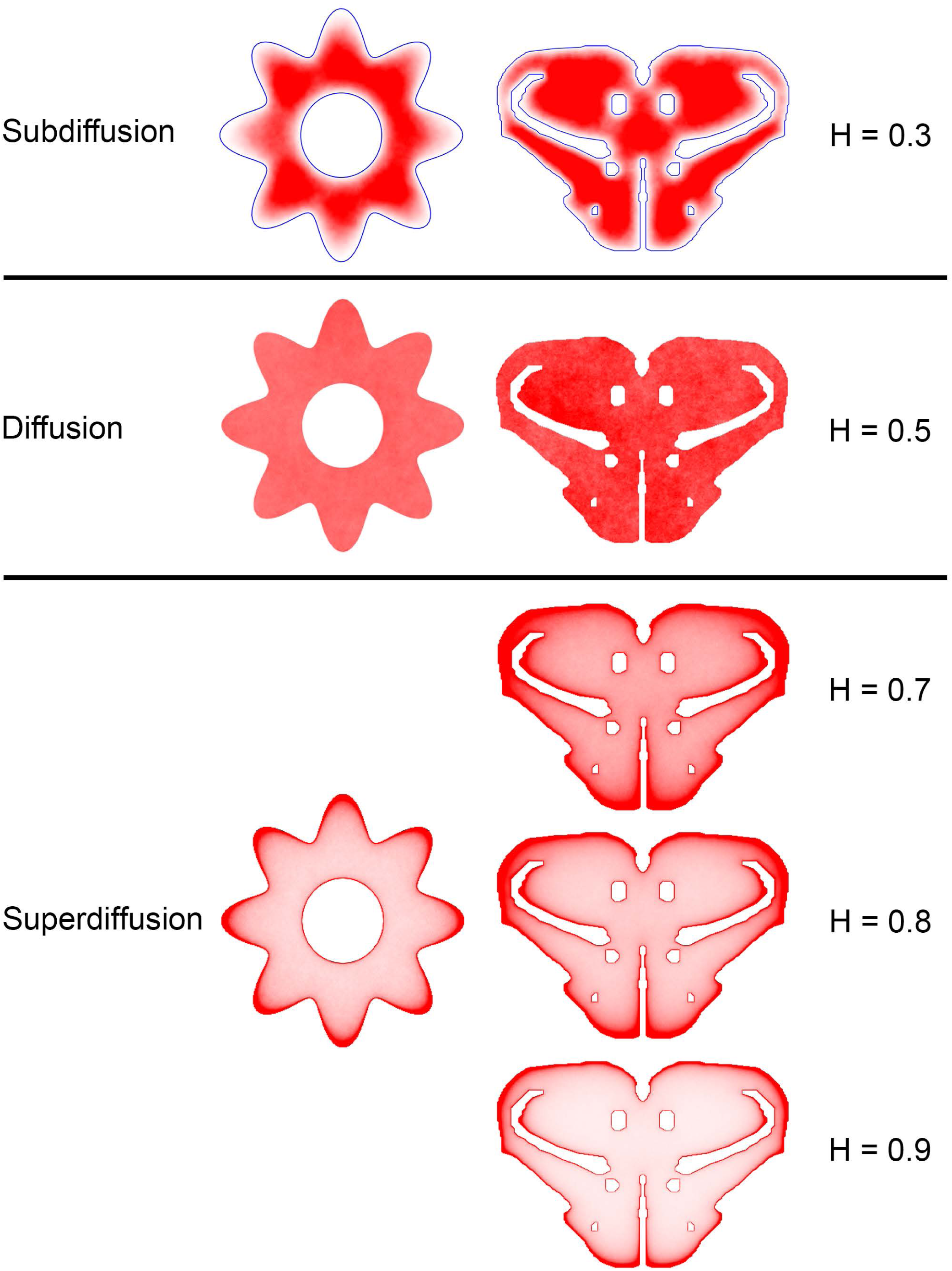
The dependence of equilibrium fiber density on the Hurst index (*H*). Two shapes were used: the cross-section of a cylinder with an outer boundary that has a non-constant curvature (representing an abstracted neural tube; left column) and a shape representing the coronal section of the mouse rostral diencephalon (also used in Figure 3; right column). The contour borders are shown in blue in the subdiffusion regime (*H* = 0.3); no borders are marked in the other regimes (but they are clearly visible due to the accumulation of fibers). The simulated densities are plotted with no transformation. One hundred fibers were used for the abstracted shape in the left column, while 960 trajectories were employed for the coronal section in the right column. The simulated densities are mapped linearly to the color scale, such that darker regions represent higher densities.

At the outer border, a considerable mismatch was observed between some *gradients* of densities (e.g., Fig. 3E). Considering the neuroanatomical simplicity of the simulated shape (e.g., it contained no “cells”), this result is not surprising. Also, the simulated gradients depend on the value of *H* and the attenuation parameter of “optical transformation” (e.g., they can be made less steep, with an effect on the overall density intensity). Matching the simulated and actual gradients precisely is difficult because the true fiber density cannot be determined in immunostained sections without tracing every fiber. Also, many non-linear effects can take place between the section and the image sensor, even at optimal illumination settings.

The strongest discrepancy was found between the strong density spikes around white matter tracts in the simulations (where the tracts were modeled as impenetrable “obstacles”) and the virtual absence of such spikes in the actual (immunostained) sections. This suggests that these tracts cannot be modeled as “hard” obstacles and that other FBM-reflection models may reflect their properties more accurately.

Since a fixed *H* was used in the simulations, we investigated the sensitivity of the obtained results to a range of *H* values (Fig. 6). The density distribution patterns varied dramatically across the three diffusion regimes, as anticipated (Wada and Vojta, 2018). However, they were robust within the superdiffusion regime, suggesting that the results can be safely generalized to other *H* values, beyond the one that was used in the simulations (0.8).

Finally, we examined the sensitivity of the results to cell packing. Heterogeneous crowding is an essential property of neural tissue (Hrabetova et al., 2018), but currently little is known about the variability of the crowding density in brain regions. In the present study, we used a relatively simple model and tested whether the accumulation of fibers at borders could be reversed by many cell-like obstacles that can potentially produce trapping effects (Fig. 7). Despite the presence of many obstacle surfaces, the simulation produced only thin, high-density layers around the obstacles, with no major effect on the overall density distribution. This result supports the robustness of our findings and is generally consistent with experimental observations in the cerebral cortex, where the high density of serotonergic fibers may not be restricted to cortical layer I (which is virtually devoid of neuron somata) and may also include layers II-III (which contain densely packed neurons) (Voigt and de Lima, 1991b). However, this finding should not be overgeneralized without future studies of the interactions among different shape geometries, heterogeneous crowding, and the physical constraints of the serotonergic fibers (e.g., the simulated trajectories in our study had no physical width).

**Figure 7.**
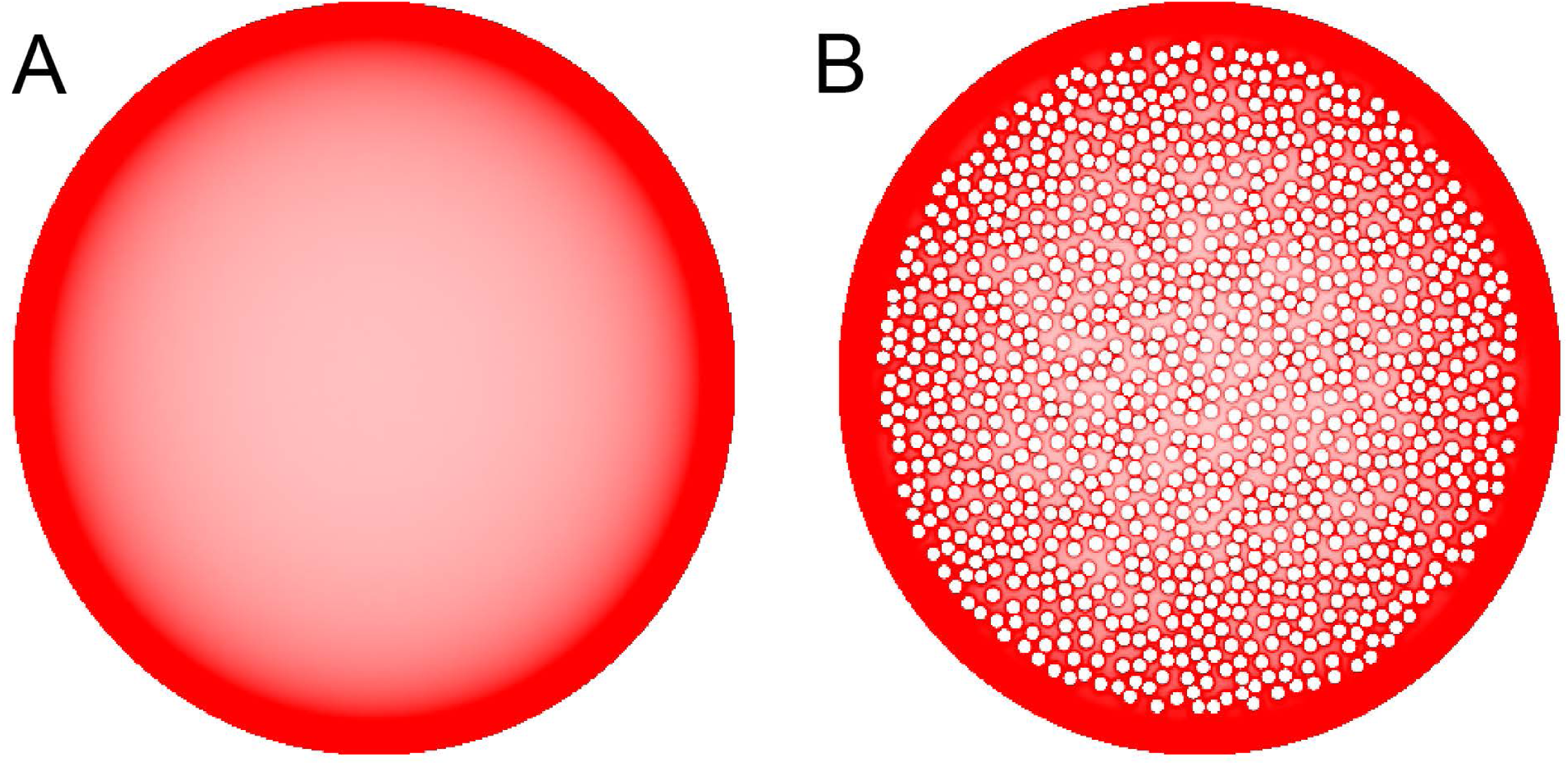
The sensitivity of equilibrium fiber density to densely packed internal obstacles. Even though fibers accumulate around the border of each individual obstacle, the obstacles have only a minor effect on the overall density distribution. In the simulations, *H* = 0.8 and 192 fibers were used. The simulated densities are mapped linearly to the color scale, such that darker regions represent higher densities.

## 4. DISCUSSION

We introduce a novel approach to the self-organization of serotonergic fibers in the brain and demonstrate that rFBM can replicate the behavior of these fibers at “hard” borders (such as the pial and ependymal surfaces of the brain). In contrast, major white matter tracts do not appear to significantly “reflect” serotonergic fibers, even though they constrain their trajectories as obstacles. This phenomenon may be due to the fact that many individual fibers can penetrate (and perhaps traverse) these tracts (Fig. 1F). The presence of serotonergic fibers in some major pathways, such as the fasciculus retroflexus, has been noted in early neuroanatomical studies (Lidov and Molliver, 1982). These phenomena can be included in our model by treating the boundaries of white matter tracts differently from other boundaries, such as modelling them as soft repulsive potentials (Vojta et al., unpublished).

Our model does not include the rich brain architecture and biological signals that may act on serotonergic fibers. While the simulated fiber densities are a good approximation of the actual fiber densities, some of these factors may be important for more accurate predictions. We briefly mention some of them.

We used two-dimensional shapes, but the brain is a three-dimensional object. This suggests that some discrepancies between the actual and simulated densities may be due to the geometry of coronal sections just rostral or caudal to the selected section, with some fibers “spilling into” or “leaking from” from the examined section. This problem may be particularly significant where the geometry of ventricular spaces rapidly changes in the rostro-caudal direction (e.g., ventricular spaces “fuse” or “separate” in two-dimensional projections). Conceptually, the computational approach can be easily extended to three dimensions, provided a fully three-dimensional model of the brain geometry is available. However, the numerical effort for such simulations would be significantly higher and may exceed the currently available numerical capabilities. One potential solution is using two-dimensional brain shapes obtained in two or three perpendicular planes (e.g., coronal, sagittal, axial), which may allow a more accurate reconstruction of densities in all three dimensions.

The brain can be viewed as a highly heterogeneous material. It is densely packed with cells, their processes, microvasculature, and other elements that serotonergic fibers cannot penetrate. Axons can be assembled into major white matter tracts which may require detours; for example, the number of axons in the primate corpus callosum can be on the order of 10^7^-10^8^ (Doty, 2007). The fine structure of the extracellular space (ECS) in the brain requires state-of-the-art experimental methods and currently is an active area of research (Nicholson and Hrabetova, 2017; Hrabetova et al., 2018). The stochastic geometry of the ECS may impose a particular covariance structure on traveling fibers (or a “memory effect”) (Morgado et al., 2002; Bénichou et al., 2013), possibly in a region-specific manner. Some of this geometry can be modeled with sphere packing (to represent cell bodies) (Picka, 2007). Similar problems arise in intracellular environments (Smith et al., 2017). In addition to the “hard” geometry, the stochastic properties of fibers may be affected by the viscoelastic properties of their environment (Cherstvy et al., 2019). In the case of FBM, it may lead to different *H* values in different brain regions, with implications for local fiber densities. Conversely, the estimated *H* values of fibers can be potentially used to obtain information about the structure and integrity of the ECS, with possible biomedical applications.

Since individual serotonergic fibers can be visualized, estimates of *H* can be obtained from experimental data. However, it requires overcoming a few technical challenges, which are a focus of our larger research program (in the present study, we assumed *H* = 0.8 and demonstrated the robustness of the main results). For example, tracing a single fiber in immunostained brain sections (e.g., 40 µm in thickness) is difficult because most fibers exit the section in the Z-direction, before advancing substantially in the X-Y plane (Janusonis et al., 2019). Modern tissue clearing methods and light-sheet microscopy allow direct 3D-imaging with no sectioning (Mano et al., 2018; Hillman et al., 2019), but light-sheet microscopy is only now approaching the sub-micrometer resolution needed for the imaging of individual serotonergic fibers (Chakraborty et al., 2019). The high fiber densities in most brain regions presents another problem, where single-fiber tracing has to be performed in the presence of interfering signals, such as other fibers and potential branching points. Advances in computer image analysis (Kayasandik et al., 2018; Falk et al., 2019), combined with transgenic technologies that allow labeling individual neurons and their processes with unique combinations of fluorophores (such as Brainbow 3.2 (Cai et al., 2013)), are well positioned to advance these efforts.

Our model assumes no self-avoidance or biological feedback signals that depend on fiber density. Several factors have been reported to control the growth of distribution of serotonergic fibers, but this research has been heavily influenced by the notion that orderly distribution cannot be achieved without tight biological control (see, for example, Chen et al. (2017)). The simulation results provide evidence that a considerable degree of self-organization can be achieved with simple assumptions, but it does not rule out these factors. Early studies have found that S100β, a biologically active protein, may promote the development of serotonergic fibers. Interestingly, S100β can be released from astrocytes, in response of activation of serotonin 5-HT_1A_ receptors, suggesting a positive feedback loop (Whitaker-Azmitia, 2001). Also, the absence of brain serotonin synthesis alters the development of normal fiber densities, but this effect appears to be strongly region-dependent (Migliarini et al., 2013). It may be mediated by the brain-derived neurotrophic factor (BDNF) which has long been implicated in the growth of serotonergic fibers (Mamounas et al., 1995; Migliarini et al., 2013). It remains unclear how serotonin affects fiber densities under physiological conditions because a different genetic model (with a less severe reduction of brain serotonin levels) has failed to reproduce these effects (Donovan et al., 2019). Recently, protocadherin-αc2 has been strongly implicated in the distribution of serotonergic fibers, through homophilic interaction between individual axons (Katori et al., 2009; Chen et al., 2017; Katori et al., 2017). Intriguingly, protocadherin-α mutants show pronounced increases in the fiber densities in layer I of the primary motor cortex and in the lacunosum-moleculare layer of the hippocampus, both of which are at the border of their respective brain regions. This study also has noted that in some other regions the “distribution of serotonin axonal terminals was […] dense at the periphery of each region but sparse in the center” (Katori et al., 2009). Our results suggest that this experimental result may reflect either a more pronounced rFBM behavior (in the absence of fiber interaction) or an rFBM with a higher *H* (induced by the genetic mutation).

It is important to note that FBM is not the only mathematical model that allows superdiffusion (or anomalous diffusion, more generally). Superdiffusion can also be modeled with Lévy flights or with continuous-time random walks (CTRWs) that have a heavy-tailed displacement probability density (Codling et al., 2008; Schulz et al., 2013; Metzler et al., 2014). These two processes allow large, instantaneous (spatially discontinuous) jumps from one location to another, but CTRWs are additionally parametrized by the waiting-time probability density (therefore, a Lévy flight can be viewed as a special CTRW). They are highly appropriate for some physical processes (e.g., the dynamics of molecular complexes jumping from one segment of a polymer to another, facilitated by folding-induced physical proximity (Lomholt et al., 2005)), but they do not realistically represent axon growth that progresses more smoothly. Also, the trajectories of growing axons are likely to show long-range temporal correlations that are an inherent property of FBM (in addition to other useful properties reviewed in the Introduction; here we assume *H ≠ 1/*2). It should be noted that the long-range correlations in FBM extend to arbitrarily large distances (Biagini et al., 2010), which may exceed biological reality, but a possible theoretical refinement may be provided by stochastic processes in which the long-range correlations are cut off at a large but finite distance (Molina-Garcia et al., 2018). In addition, branching FBM-like processes may offer insights into how the bifurcation or arborization of serotonergic fibers can affect their steady state distribution. Direct simulations of macromolecular dynamics in confined domains can further enrich these studies; for example, in some simulations particles near the wall tend to stay near the wall (Chow and Skolnick, 2015), which may explain the tendency of serotonergic fibers to orient parallel to the edge immediately below the pia (for depths up to 25-50 µm; Fig. 1). Finally, it has been recently demonstrated (Vojta et al., 2019) that the increased density close to a boundary arises from the nonequilibrium nature of FBM. A similar anomalous diffusion process in thermal equilibrium, modelled by the fractional Langevin equation, does not lead to accumulation at the boundaries. This property of FBM is consistent with the active growth of the serotonergic fibers.

In summary, the present study demonstrates that FBM offers a promising theoretical framework for the modeling of serotonergic fibers. Since serotonin-releasing fibers are a part of the larger ascending reticular activating system, which releases other major neurotransmitters and has widespread projections, this framework may also be useful in advancing the understanding of other stochastic axon systems in vertebrate brains.

## ACKNOWLEDGEMENTS

We thank Kasie Mays (UCSB) for the assistance with tissue preparation.

## AUTHOR CONTRIBUTIONS

SJ proposed the hypothesis of serotonergic fibers as paths of a stochastic process, performed the immunostaining, prepared the 2D-shape matrices for simulations, and wrote the first draft of the manuscript. TV guided the computational analyses of reflected FBM and performed all supercomputing simulations. ND suggested FBM as a potential model that allows scale-invariance and made other theoretical contributions. RM led the development of the model within the theoretical framework of anomalous diffusion processes. All authors are the Principal Investigators of their respective research programs.

## CONTRIBUTION TO THE FIELD

This study is the first application of Fractional Brownian Motion to the investigation of serotonergic fibers in the brain. In a novel approach, it uses supercomputing simulations to bridge experimental neurobiology and recent findings in reflected Fractional Brownian Motion. The present work is a collaboration among researchers in all of these fields and will motivate further advances in both the neurobiology of the ascending reticular activating system and the theory of stochastic processes.

## REFERENCES

Agnati, L.F., and Fuxe, K. (2014). Extracellular-vesicle type of volume transmission and tunnelling-nanotube type of wiring transmission add a new dimension to brain neuro-glial networks. Philos Trans R Soc Lond B Biol Sci 369(1652). doi: 10.1098/rstb.2013.0505.

Azmitia, E.C., Singh, J.S., and Whitaker-Azmitia, P.M. (2011). Increased serotonin axons (immunoreactive to 5-HT transporter) in postmortem brains from young autism donors. Neuropharmacology 60(7-8), 1347–1354. doi: 10.1016/j.neuropharm.2011.02.002.

Bénichou, O., Bodrova, A., Chakraborty, D., Illien, P., Law, A., Mejía-Monasterio, C., et al. (2013). Geometry-Induced Superdiffusion in Driven Crowded Systems. Physical Review Letters 111.

Benzekhroufa, K., Liu, B., Tang, F., Teschemacher, A.G., and Kasparov, S. (2009). Adenoviral vectors for highly selective gene expression in central serotonergic neurons reveal quantal characteristics of serotonin release in the rat brain. BMC Biotechnol 9, 23. doi: 10.1186/1472-6750-9-23.

Biagini, F., Hu, Y., Oksendal, B., and Zhang, T. (2010). Stochastic Calculus for Fractional Brownian Motion and Applications. London: Springer.

Cai, D., Cohen, K.B., Luo, T., Lichtman, J.W., and Sanes, J.R. (2013). Improved tools for the Brainbow toolbox. Nat Methods 10(6), 540–547.

Cardozo Pinto, D.F., Yang, H., Pollak Dorocic, I., de Jong, J.W., Han, V.J., Peck, J.R., et al. (2019). Characterization of transgenic mouse models targeting neuromodulatory systems reveals organizational principles of the dorsal raphe. Nat Commun 10(1), 4633. doi: 10.1038/s41467-019-12392-2.

Carrera, I., Molist, P., Anadon, R., and Rodriguez-Moldes, I. (2008). Development of the serotoninergic system in the central nervous system of a shark, the lesser spotted dogfish Scyliorhinus canicula. J Comp Neurol 511(6), 804–831. doi: 10.1002/cne.21857.

Chakraborty, T., Driscoll, M.K., Jeffery, E., Murphy, M.M., Roudot, P., Chang, B.J., et al. (2019). Light-sheet microscopy of cleared tissues with isotropic, subcellular resolution. Nat Methods 16(11), 1109–1113. doi: 10.1038/s41592-019-0615-4.

Chen, W.V., Nwakeze, C.L., Denny, C.A., O’Keeffe, S., Rieger, M.A., Mountoufaris, G., et al. (2017). Pcdhalphac2 is required for axonal tiling and assembly of serotonergic circuitries in mice. Science 356(6336), 406–411. doi: 10.1126/science.aal3231.

Cherstvy, A.G., Thapa, S., Wagner, C.E., and Metzler, R. (2019). Non-Gaussian, non-ergodic, and non-Fickian diffusion of tracers in mucin hydrogels. Soft Matter 15(12), 2526–2551. doi: 10.1039/c8sm02096e.

Chow, E., and Skolnick, J. (2015). Effects of confinement on models of intracellular macromolecular dynamics. Proceedings of the National Academy of Science 112, 14846–14851.

Codling, E.A., Plank, M.J., and Benhamou, S. (2008). Random walk models in biology. J R Soc Interface 5(25), 813–834. doi: 10.1098/rsif.2008.0014.

Donovan, L.J., Spencer, W.C., Kitt, M.M., Eastman, B.A., Lobur, K.J., Jiao, K., et al. (2019). Lmx1b is required at multiple stages to build expansive serotonergic axon architectures. Elife 8. doi: 10.7554/eLife.48788.

Doty, R.W. (2007). “Cortical Commissural Connections in Primates,” in Evolution of Nervous Systems: A Comprehensive Reference, eds. J.H. Kaas & T.M. Preuss. (New York: Academic Press), 277–289.

Dougherty, S.E., Kajstura, T.J., Jin, Y., Chan-Cortes, M.H., Kota, A., and Linden, D.J. (2019). Catecholaminergic axons in the neocortex of adult mice regrow following brain injury. Exp Neurol 323, 113089. doi: 10.1016/j.expneurol.2019.113089.

Falk, T., Mai, D., Bensch, R., Cicek, O., Abdulkadir, A., Marrakchi, Y., et al. (2019). U-Net: deep learning for cell counting, detection, and morphometry. Nat Methods 16(1), 67–70. doi: 10.1038/s41592-018-0261-2.

Gagnon, D., and Parent, M. (2014). Distribution of VGLUT3 in highly collateralized axons from the rat dorsal raphe nucleus as revealed by single-neuron reconstructions. PLoS One 9(2), e87709. doi: 10.1371/journal.pone.0087709.

Guggenberger, T., Pagnini, G., Vojta, T., and Metzler, R. (2019). Fractional Brownian motion in a finite interval: correlations effect depletion or accretion zones of particles near boundaries. New Journal of Physics 21(022002).

Hawthorne, A.L., Hu, H., Kundu, B., Steinmetz, M.P., Wylie, C.J., Deneris, E.S., et al. (2011). The unusual response of serotonergic neurons after CNS Injury: lack of axonal dieback and enhanced sprouting within the inhibitory environment of the glial scar. J Neurosci 31(15), 5605–5616. doi: 10.1523/jneurosci.6663-10.2011.

Hawthorne, A.L., Wylie, C.J., Landmesser, L.T., Deneris, E.S., and Silver, J. (2010). Serotonergic neurons migrate radially through the neuroepithelium by dynamin-mediated somal translocation. J Neurosci 30(2), 420–430. doi: 10.1523/jneurosci.2333-09.2010.

Hendricks, T., Francis, N., Fyodorov, D., and Deneris, E.S. (1999). The ETS domain factor Pet-1 is an early and precise marker of central serotonin neurons and interacts with a conserved element in serotonergic genes. J Neurosci 19(23), 10348–10356.

Hillman, E.M.C., Voleti, V., Li, W., and Yu, H. (2019). Light-Sheet Microscopy in Neuroscience. Annu Rev Neurosci 42, 295–313. doi: 10.1146/annurev-neuro-070918-050357.

Hornung, J.P. (2003). The human raphe nuclei and the serotonergic system. J Chem Neuroanat 26(4), 331–343. doi: 10.1016/j.jchemneu.2003.10.002.

Hrabetova, S., Cognet, L., Rusakov, D.A., and Nagerl, U.V. (2018). Unveiling the Extracellular Space of the Brain: From Super-resolved Microstructure to In Vivo Function. J Neurosci 38(44), 9355–9363. doi: 10.1523/jneurosci.1664-18.2018.

Ito, K., and McKean, H.P. (1965). Diffusion Processes and Their Sample Paths.

Ivgy-May, N., Tamir, H., and Gershon, M.D. (1994). Synaptic properties of serotonergic growth cones in developing rat brain. J Neurosci 14(3 Pt 1), 1011–1029.

Jacobs, B.L., and Azmitia, E.C. (1992). Structure and function of the brain serotonin system. Physiol Rev 72(1), 165–229. doi: 10.1152/physrev.1992.72.1.165.

Janusonis, S., and Detering, N. (2019). A stochastic approach to serotonergic fibers in mental disorders. Biochimie 161, 15–22. doi: 10.1016/j.biochi.2018.07.014.

Janusonis, S., Gluncic, V., and Rakic, P. (2004). Early serotonergic projections to Cajal-Retzius cells: relevance for cortical development. J Neurosci 24(7), 1652–1659. doi: 10.1523/jneurosci.4651-03.2004.

Janusonis, S., Mays, K.C., and Hingorani, M.T. (2019). Serotonergic Axons as 3D-Walks. ACS Chem Neurosci 10(7), 3064–3067. doi: 10.1021/acschemneuro.8b00667.

Jin, Y., Dougherty, S.E., Wood, K., Sun, L., Cudmore, R.H., Abdalla, A., et al. (2016). Regrowth of Serotonin Axons in the Adult Mouse Brain Following Injury. Neuron 91(4), 748–762. doi: 10.1016/j.neuron.2016.07.024.

Kajstura, T.J., Dougherty, S.E., and Linden, D.J. (2018). Serotonin axons in the neocortex of the adult female mouse regrow after traumatic brain injury. J Neurosci Res 96(4), 512–526. doi: 10.1002/jnr.24059.

Katori, S., Hamada, S., Noguchi, Y., Fukuda, E., Yamamoto, T., Yamamoto, H., et al. (2009). Protocadherin-alpha family is required for serotonergic projections to appropriately innervate target brain areas. J Neurosci 29(29), 9137–9147. doi: 10.1523/jneurosci.5478-08.2009.

Katori, S., Noguchi-Katori, Y., Okayama, A., Kawamura, Y., Luo, W., Sakimura, K., et al. (2017). Protocadherin-alphaC2 is required for diffuse projections of serotonergic axons. Sci Rep 7(1), 15908. doi: 10.1038/s41598-017-16120-y.

Kayasandik, C., Negi, P., Laezza, F., Papadakis, M., and Labate, D. (2018). Automated sorting of neuronal trees in fluorescent images of neuronal networks using NeuroTreeTracer. Sci Rep 8(1), 6450. doi: 10.1038/s41598-018-24753-w.

Kiyasova, V., and Gaspar, P. (2011). Development of raphe serotonin neurons from specification to guidance. Eur J Neurosci 34(10), 1553–1562. doi: 10.1111/j.1460-9568.2011.07910.x.

Lidov, H.G., and Molliver, M.E. (1982). An immunohistochemical study of serotonin neuron development in the rat: ascending pathways and terminal fields. Brain Res Bull 8(4), 389–430. doi: 10.1016/0361-9230(82)90077-6.

Linley, S.B., Hoover, W.B., and Vertes, R.P. (2013). Pattern of distribution of serotonergic fibers to the orbitomedial and insular cortex in the rat. J Chem Neuroanat 48-49, 29–45. doi: 10.1016/j.jchemneu.2012.12.006.

Lomholt, M.A., Ambjornsson, T., and Metzler, R. (2005). Optimal target search on a fast-folding polymer chain with volume exchange. Phys Rev Lett 95(26), 260603. doi: 10.1103/PhysRevLett.95.260603.

Mai, J.K., and Ashwell, K.W.S. (2004). “Fetal Development of the Central Nervous System,” in The Human Nervous System, eds. G. Paxinos & J.K. Mai. Elsevier), 49–94.

Maia, G.H., Soares, J.I., Almeida, S.G., Leite, J.M., Baptista, H.X., Lukoyanova, A.N., et al. (2019). Altered serotonin innervation in the rat epileptic brain. Brain Res Bull 152, 95–106. doi: 10.1016/j.brainresbull.2019.07.009.

Makse, H.A., Havlin, S., Schwartz, M., and Stanley, H.E. (1996). Method for generating long-range correlations for large systems. Physical Review E 53, 5445–5449.

Mamounas, L.A., Blue, M.E., Siuciak, J.A., and Altar, C.A. (1995). Brain-derived neurotrophic factor promotes the survival and sprouting of serotonergic axons in rat brain. J Neurosci 15(12), 7929–7939.

Mandelbrot, B.B., and Van Ness, J.W. (1968). Fractional Brownian motions, fractional noises and applications. SIAM Review 10, 422–437.

Mano, T., Albanese, A., Dodt, H.U., Erturk, A., Gradinaru, V., Treweek, J.B., et al. (2018). Whole-Brain Analysis of Cells and Circuits by Tissue Clearing and Light-Sheet Microscopy. J Neurosci 38(44), 9330–9337. doi: 10.1523/jneurosci.1677-18.2018.

Metzler, R., Jeon, J.H., Cherstvy, A.G., and Barkai, E. (2014). Anomalous diffusion models and their properties: non-stationarity, non-ergodicity, and ageing at the centenary of single particle tracking. Phys Chem Chem Phys 16(44), 24128–24164. doi: 10.1039/c4cp03465a.

Migliarini, S., Pacini, G., Pelosi, B., Lunardi, G., and Pasqualetti, M. (2013). Lack of brain serotonin affects postnatal development and serotonergic neuronal circuitry formation. Mol Psychiatry 18(10), 1106–1118. doi: 10.1038/mp.2012.128.

Molina-Garcia, D., Sandev, T., Safdari, H., Pagnini, G., Chechkin, A., and Metzler, R. 2018. Crossover from anomalous to normal diffusion: truncated power-law noise correlations and applications to dynamics in lipid bilayers. arXiv e-prints [Online]. Available: https://ui.adsabs.harvard.edu/abs/2018arXiv180909586M [Accessed September 01, 2018].

Morgado, R., Oliveira, F.A., Batrouni, G.G., and Hansen, A. (2002). Relation between anomalous and normal diffusion in systems with memory. Phys Rev Lett 89(10), 100601. doi: 10.1103/PhysRevLett.89.100601.

Morin, L.P., and Meyer-Bernstein, E.L. (1999). The ascending serotonergic system in the hamster: comparison with projections of the dorsal and median raphe nuclei. Neuroscience 91(1), 81–105. doi: 10.1016/s0306-4522(98)00585-5.

Mosienko, V., Beis, D., Pasqualetti, M., Waider, J., Matthes, S., Qadri, F., et al. (2015). Life without brain serotonin: reevaluation of serotonin function with mice deficient in brain serotonin synthesis. Behav Brain Res 277, 78–88. doi: 10.1016/j.bbr.2014.06.005.

Nicholson, C., and Hrabetova, S. (2017). Brain Extracellular Space: The Final Frontier of Neuroscience. Biophys J 113(10), 2133–2142. doi: 10.1016/j.bpj.2017.06.052.

Numasawa, Y., Hattori, T., Ishiai, S., Kobayashi, Z., Kamata, T., Kotera, M., et al. (2017). Depressive disorder may be associated with raphe nuclei lesions in patients with brainstem infarction. J Affect Disord 213, 191–198. doi: 10.1016/j.jad.2017.02.005.

Okaty, B.W., Commons, K.G., and Dymecki, S.M. (2019). Embracing diversity in the 5-HT neuronal system. Nat Rev Neurosci 20(7), 397–424. doi: 10.1038/s41583-019-0151-3.

Papadopoulos, G.C., Parnavelas, J.G., and Buijs, R. (1987). Monoaminergic fibers form conventional synapses in the cerebral cortex. Neurosci Lett 76(3), 275–279. doi: 10.1016/0304-3940(87)90414-9.

Picka, J. 2007. Statistical Inference for Disordered Sphere Packings. arXiv e-prints [Online]. Available: https://ui.adsabs.harvard.edu/abs/2007arXiv0711.3035P [Accessed November 01, 2007].

Polovnikov, K.E., Gherardi, M., Cosentino-Lagomarsino, M., and Tamm, M.V. (2018). Fractal Folding and Medium Viscoelasticity Contribute Jointly to Chromosome Dynamics. Physical Review Letters 120.

Polovnikov, K.E., Nechaev, S., and Tamm, M.V. (2019). Many-body contacts in fractal polymer chains and fractional Brownian trajectories. Physical Review E 99.

Pratelli, M., Migliarini, S., Pelosi, B., Napolitano, F., Usiello, A., and Pasqualetti, M. (2017). Perturbation of Serotonin Homeostasis during Adulthood Affects Serotonergic Neuronal Circuitry. eNeuro 4(2). doi: 10.1523/eneuro.0376-16.2017.

Pratelli, M., and Pasqualetti, M. (2019). Serotonergic neurotransmission manipulation for the understanding of brain development and function: Learning from Tph2 genetic models. Biochimie 161, 3–14. doi: 10.1016/j.biochi.2018.11.016.

Qian, H. (2003). “Fractional Brownian motion and fractional Gaussian noise,” in Processes with Long-Range Correlations: Theory and Applications, eds. G. Rangarajan & M. Ding. (Berlin/Heidelberg: Springer), 22–33.

Quentin, E., Belmer, A., and Maroteaux, L. (2018). Somato-Dendritic Regulation of Raphe Serotonin Neurons; A Key to Antidepressant Action. Front Neurosci 12, 982. doi: 10.3389/fnins.2018.00982.

Ren, J., Friedmann, D., Xiong, J., Liu, C.D., Ferguson, B.R., Weerakkody, T., et al. (2018). Anatomically Defined and Functionally Distinct Dorsal Raphe Serotonin Sub-systems. Cell 175(2), 472-487.e420. doi: 10.1016/j.cell.2018.07.043.

Schneeberger, M., Parolari, L., Das Banerjee, T., Bhave, V., Wang, P., Patel, B., et al. (2019). Regulation of Energy Expenditure by Brainstem GABA Neurons. Cell 178(3), 672-685.e612. doi: 10.1016/j.cell.2019.05.048.

Schulz, J.H.P., Chechkin, A.V., and Metzler, R. (2013). Correlated continuous time random walks: combining scale-invariance with long-range memory for spatial and temporal dynamics. Journal of Physics A Mathematical General 46.

Slaten, E.R., Hernandez, M.C., Albay, R., 3rd, Lavian, R., and Janusonis, S. (2010). Transient expression of serotonin 5-HT4 receptors in the mouse developing thalamocortical projections. Dev Neurobiol 70(3), 165–181. doi: 10.1002/dneu.20775.

Smith, S., Cianci, C., and Grima, R. (2017). Macromolecular crowding directs the motion of small molecules inside cells. J R Soc Interface 14(131). doi: 10.1098/rsif.2017.0047.

Soiza-Reilly, M., Anderson, W.B., Vaughan, C.W., and Commons, K.G. (2013). Presynaptic gating of excitation in the dorsal raphe nucleus by GABA. Proc Natl Acad Sci U S A 110(39), 15800–15805. doi: 10.1073/pnas.1304505110.

Stamp, J.A., and Semba, K. (1995). Extent of colocalization of serotonin and GABA in the neurons of the rat raphe nuclei. Brain Res 677(1), 39–49. doi: 10.1016/0006-8993(95)00119-b.

Steinbusch, H.W. (1981). Distribution of serotonin-immunoreactivity in the central nervous system of the rat-cell bodies and terminals. Neuroscience 6(4), 557–618. doi: 10.1016/0306-4522(81)90146-9.

Stuesse, S.L., Cruce, W.L., and Northcutt, R.G. (1991). Localization of serotonin, tyrosine hydroxylase, and leu-enkephalin immunoreactive cells in the brainstem of the horn shark, Heterodontus francisci. J Comp Neurol 308(2), 277–292. doi: 10.1002/cne.903080211.

Sundstrom, E., Kolare, S., Souverbie, F., Samuelsson, E.B., Pschera, H., Lunell, N.O., et al. (1993). Neurochemical differentiation of human bulbospinal monoaminergic neurons during the first trimester. Brain Res Dev Brain Res 75(1), 1–12. doi: 10.1016/0165-3806(93)90059-j.

Voigt, T., and de Lima, A.D. (1991a). Serotoninergic innervation of the ferret cerebral cortex. I. Adult pattern. J Comp Neurol 314(2), 403–414. doi: 10.1002/cne.903140214.

Voigt, T., and de Lima, A.D. (1991b). Serotoninergic innervation of the ferret cerebral cortex. II. Postnatal development. J Comp Neurol 314(2), 415–428. doi: 10.1002/cne.903140215.

Vojta, T., Skinner, S., and Metzler, R. (2019). Probability density of the fractional Langevin equation with reflecting walls. Phys Rev E 100(4-1), 042142. doi: 10.1103/PhysRevE.100.042142.

Wada, A.H.O., and Vojta, T. (2018). Fractional Brownian motion with a reflecting wall. Phys Rev E 97(2-1), 020102. doi: 10.1103/PhysRevE.97.020102.

Wada, A.H.O., Warhover, A., and Vojta, T. (2019). Non-Gaussian behavior of reflected fractional Brownian motion. Journal of Statistical Mechanics: Theory and Experiment 3, 033209.

Wassie, A.T., Zhao, Y., and Boyden, E.S. (2019). Expansion microscopy: principles and uses in biological research. Nat Methods 16(1), 33–41. doi: 10.1038/s41592-018-0219-4.

Whitaker-Azmitia, P.M. (2001). Serotonin and brain development: role in human developmental diseases. Brain Res Bull 56(5), 479–485. doi: 10.1016/s0361-9230(01)00615-3.

